# Natural selection shaped the rise and fall of passenger pigeon genomic diversity

**DOI:** 10.1101/154294

**Authors:** Gemma G. R. Murray, André E. R. Soares, Ben J. Novak, Nathan K. Schaefer, James A. Cahill, Allan J. Baker, John R. Demboski, Andrew Doll, Rute R. Da Fonseca, Tara L. Fulton, M. Thomas P. Gilbert, Peter D. Heintzman, Brandon Letts, George McIntosh, Brendan L. O’Connell, Mark Peck, Marie-Lorraine Pipes, Edward S. Rice, Kathryn M. Santos, A. Gregory Sohrweide, Samuel H. Vohr, Russell B. Corbett-Detig, Richard E. Green, Beth Shapiro

**Author notes:** These authors contributed equally to this work (Gemma G. R. Murray, André E. R. Soares). deceased author (Allan J. Baker).

## Abstract

The extinct passenger pigeon was once the most abundant bird in North America, and possibly the world. While theory predicts that large populations will be more genetically diverse and respond more efficiently to selection, passenger pigeon genetic diversity was surprisingly low. To investigate this we analysed 41 mitochondrial and 4 nuclear genomes from passenger pigeons, and 2 genomes from band-tailed pigeons, passenger pigeons’ closest living relatives. We find that passenger pigeons’ large population size allowed for faster adaptive evolution and removal of harmful mutations, but that this drove a huge loss in neutral genetic diversity. These results demonstrate how great an impact selection can have on a vertebrate genome, and invalidate previous results that suggested population instability contributed to this species’ surprisingly rapid extinction.

---

The passenger pigeon (*Ectopistes migratorius*) numbered between 3 and 5 billion individuals prior to its 19th century decline and eventual extinction (*1*). Passenger pigeons were highly mobile, bred in large social colonies, and their population lacked clear geographic structure (*2*). Few vertebrates have populations this large and cohesive, and according to the neutral model of molecular evolution, this should lead to high genetic diversity (*3*). Preliminary analyses of passenger pigeon genomes have, however, revealed similar genetic diversity to birds with population sizes three orders of magnitude smaller (*4*). This has been interpreted within the framework of the neutral theory of molecular evolution as the result of a history of dramatic demographic fluctuations (*4*). However, in large populations natural selection may be particularly important in shaping genetic diversity: selection on one locus can cause the loss of diversity at physically linked loci (*5*–*8*), and natural selection is predicted to be more efficient in species with larger population sizes (*9*). It has been suggested that this impact of selection on genetic diversity is widespread and that it explains the long standing paradox of population genetics that the genetic diversity of a species is poorly predicted by its population size (*3*, *10*, *11*). Here we investigate the impact of natural selection on passenger pigeon genomes through comparative genomic analyses of both passenger pigeons and one of their closest living relatives, band-tailed pigeons (*Patagioenas fasciata*) (*12*). While ecologically and physiologically similar to passenger pigeons, band-tailed pigeons have a present-day population size three orders of magnitude smaller (*2*, *13*).

We first applied a Bayesian skyline model of ancestral population dynamics to the mitochondrial genomes of 41 passenger pigeons from across their former breeding range (Fig. 1A and table S1) using a lineage-specific evolutionary rate estimate (*14*). This returned a most recent effective population size (*N_e_*) of 13 million (95% HPD: 2-58 million) and similar, stable *N_e_* for the previous 50,000 years (Fig. 1B). While this *N_e_* is much lower than the census population size, it is greater than previous estimates based on analyses of nuclear genomes (*4*), and it is likely to be conservative [see supplementary materials section 2.1 (SM2.1)].

**Fig. 1.**
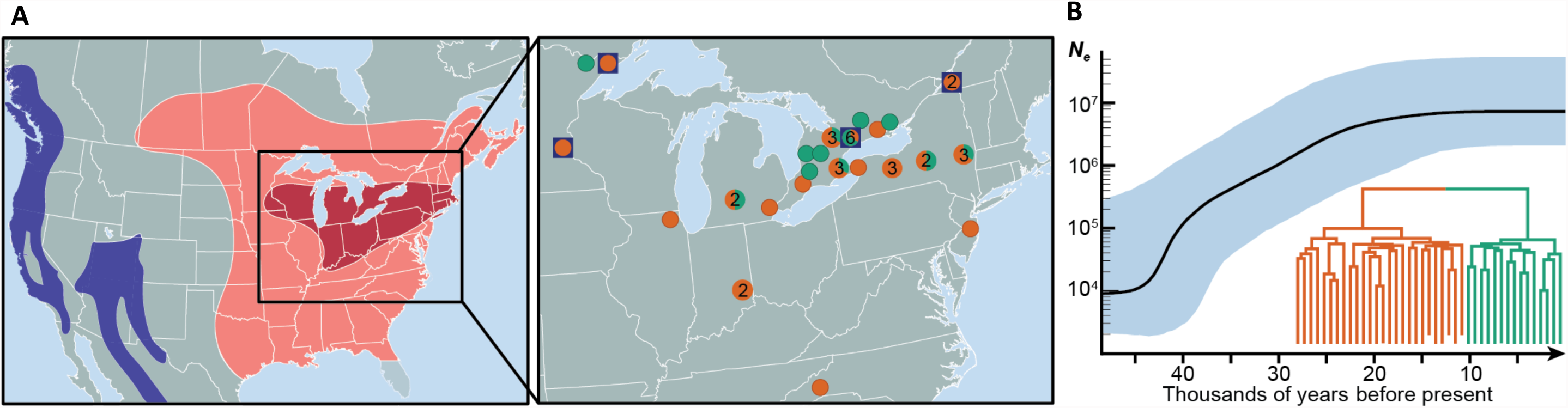
Passenger pigeon range, sample origins, and *N_e_* estimate from mitochondrial genomes. (**A**) Range of passenger pigeons at time of European contact (dark red: breeding range; light red: full range) (*1*) and current range of band-tailed pigeons (purple) (*13*), with inset showing the location of origin of the 41 passenger pigeon samples analyzed here. Locations of the four samples from which nuclear genomes were generated are indicated with a blue box. (**B**) Inferred *N_e_* and mitochondrial phylogeny from a Bayesian coalescent analysis. Colors in (**A**) inset match the phylogeny in (**B**). The structure of the phylogeny does not correlate with geography, which is consistent with an absence of geographic population structure.

We then compared nucleotide diversity (*π*) in the passenger pigeon nuclear genome to *π* in the band-tailed pigeon nuclear genome. We analysed four high-coverage passenger pigeon genome assemblies (two newly sequenced and two from published raw data; table S2), and two high-coverage band-tailed pigeon genome assemblies. We found that *π* was greater in passenger pigeons (average *π* = 0.008) than in band-tailed pigeons (average *π* = 0.004), but that the difference is less than expected given their population sizes. We estimated *π* for non-overlapping 5 Mb windows across the genome, and found that these species had a correlated regional variation in *π*, but with much greater variation in passenger pigeons (Fig. 2A and fig. S1).

**Fig. 2.**
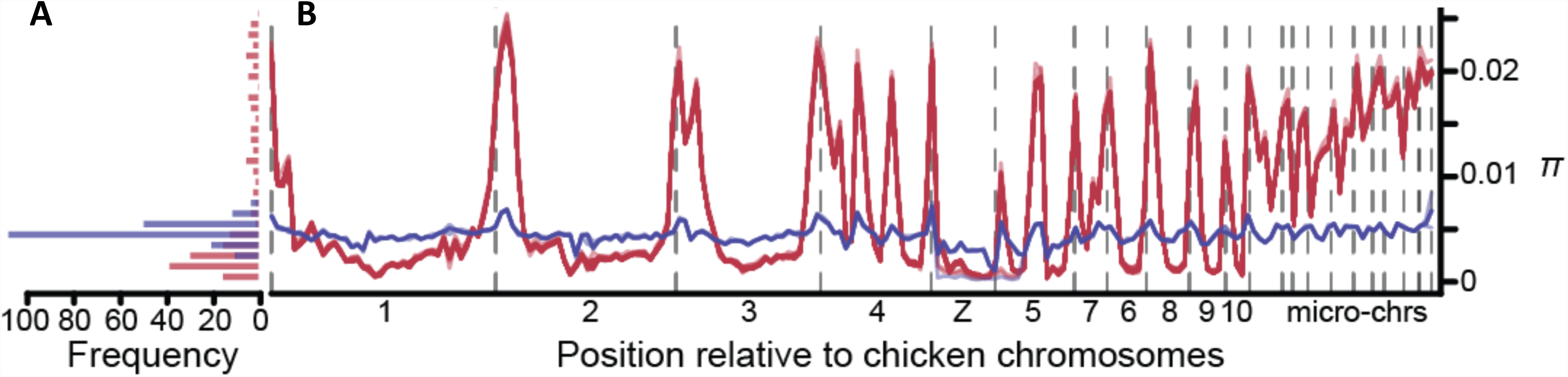
π across passenger pigeon and band-tailed pigeon genomes. (**A**) A histogram describing mean *π* for 5 Mb windows across the passenger pigeon (red) and band-tailed pigeon (blue) genomes. (**B**) Genomic distribution of individual pairwise estimates of mean *π* in 5 Mb windows across the two species’ genomes. Each between- and within-individual pairwise comparison is plotted as red (28 passenger pigeon comparisons) or blue (6 band-tailed pigeon comparisons) lines. Chromosome boundaries are indicated as vertical dashed lines. Chromosomes are ordered by their size in the chicken genome.

To explore this variation, we mapped our scaffolds to the chicken genome assembly (*15*), which should approximate the correct chromosomal structure since karyotype and synteny are strongly conserved across birds (*16*). We found that low genetic diversity regions of the passenger pigeon genome are generally in the centres of macrochromosomes, while the edges of macrochromosomes and microchromosomes have much higher diversity (Fig. 2B). Although this pattern is largely absent from the band-tailed pigeon genome, it is not likely an artefact of ancient DNA damage: our assemblies had high coverage depth (table S2), we used conservative cut-offs for calling variants, and we recovered similar patterns after excluding variants more likely to result from damage (fig. S2; SM2.2).

We next investigated the impact of selection on the evolution of protein-coding regions of the genome in both species. We calculated the rate of adaptive substitution relative to the rate of neutral substitution (*ω_a_*) and the ratio of nonsynonymous to synonymous polymorphism (*pN*/*pS*) for 5 Mb windows across the genome. A higher *ω_a_* suggests stronger or more efficient positive selection, and a lower *pN*/*pS* suggests stronger or more efficient selective constraint. We found that *ω_a_* was higher (Mann-Whitney U test, *p* = 1.1x10^−4^) and *pN*/*pS* lower (*p* = 2.7x10^−12^) in passenger pigeons than band-tailed pigeons (Fig. 3 and fig. S3). We also found that *ω_a_* was higher (*p* = 1.2x10^−8^) and *pN*/*pS* lower (*p* = 6.6x10^−10^) in high-diversity regions of the passenger pigeon genome compared to low-diversity regions (Fig. 3 and fig. S3). In addition, we found that codon usage bias, which is thought to reflect selection for translational optimization (*17*), was greater in passenger pigeons than in band-tailed pigeons, and greater in high-diversity regions (SM2.3).

**Fig. 3.**
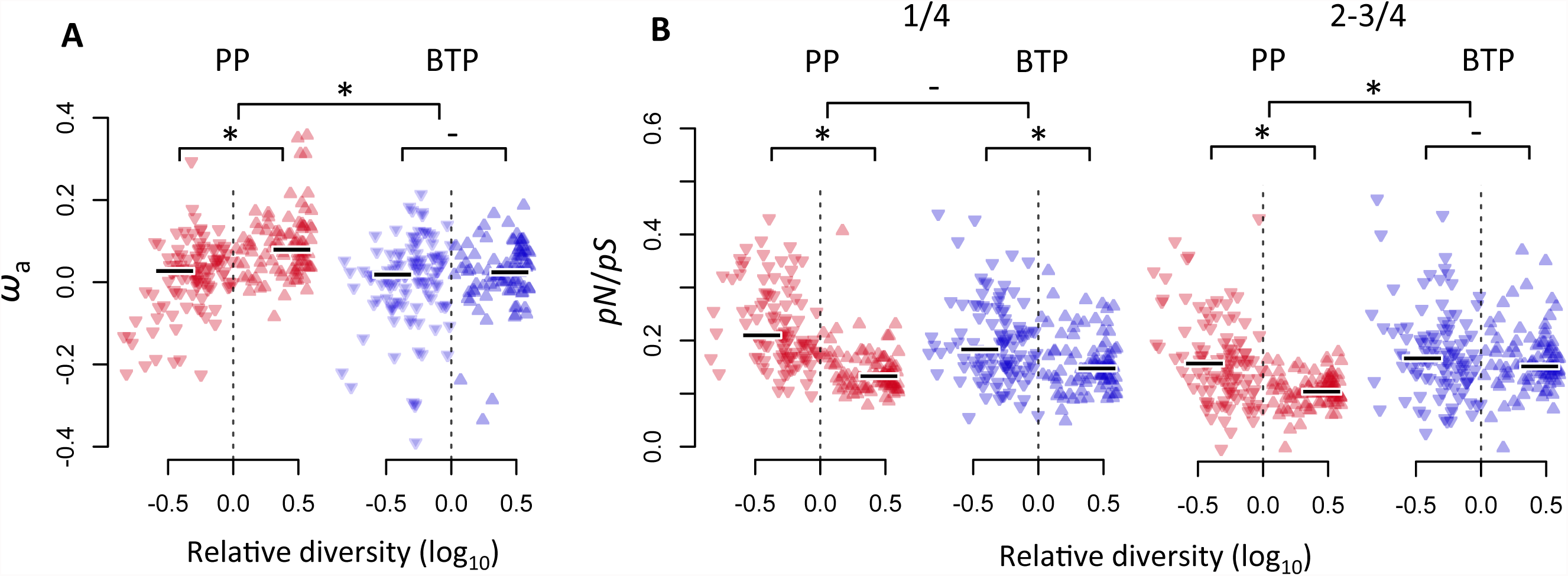
Estimates of *ω_a_* and *pN/pS*. Estimates are averages for 5 Mb windows and are plotted against the window’s genetic diversity in passenger pigeons relative to band-tailed pigeons (on a log_10_-scale). Comparisons are drawn between (**A**) *ω_a_* and (**B**) *pN*/*pS* in passenger pigeons (PP; red) and band-tailed pigeons (BTP; blue), and between low-diversity (*π*_PP_ < *π*_BTP_; point-down triangles) and high-diversity (*π*_PP_ > *π*_BTP_; point-up triangles) windows (median values are shown as horizontal lines; ‘*’ indicates *p* ≤ 1x10^−4^ and ‘-’*p* ≥ 0.1 in a Mann-Whitney U test). In (**B**) *pN*/*pS* estimates are for derived mutations present in 1/4 and 2-3/4 individuals. A higher *pN*/*pS* for lower frequency mutations could reflect the slow purging of weakly deleterious mutations. Estimates are based on analyses of two individuals from each species (see figure S3 for estimates using all passenger pigeon samples).

We also estimated the difference between the proportions of substitutions and polymorphisms that are nonsynonymous (the direction of selection, DoS) for individual genes, where a positive DoS indicates adaptive evolution. We found that DoS was more often positive in passenger pigeons than in band-tailed pigeons and, in passenger pigeons, DoS was correlated with diversity (fig. S4). McDonald-Kreitman tests identified 32 genes with strong evidence of adaptive evolution in passenger pigeons (table S3). Among them are genes associated with immune defense (e.g. *CPD (18)*), seasonal consumption of high-sugar foods in passerine birds (*SI*) (*19*), and stress modulation (*FAAH*) (*20*). Selection on these gene functions is consistent with the ecology of passenger pigeons: they had a distinctive diet (*2*), and larger and denser populations tend to endure an increased burden of transmissible pathogens (*21*) and social stress (*22*).

Differences in the efficacy of selection between passenger pigeons and band-tailed pigeons could derive from several factors (e.g. recombination rate, mutation rate, the distribution of fitness effects). However, since the close relationship between these species makes substantial differences in most of these factors unlikely, the most parsimonious explanation is the difference in population size. Theory predicts that larger populations will experience a greater efficacy of natural selection, and evidence of this has been found in comparisons across a number of other species (*9*, cf. *23*).

A greater efficacy of selection could lead to a greater impact of selection on linked sites: selection can lead to both reduced diversity at linked neutral sites and reduced efficacy of selection at linked selected sites (*3*, *5*–*8*, *24*). The impact of this will be greater where recombination rates are low, and in bird genomes recombination rates are lower in the centers of macrochromosomes, relative both to their edges and to the microchromosomes (*16*) (SM2.4). Therefore, the recombination landscape of the bird genome, combined with the an extremely large population size, could have driven the patterns we observe across the passenger pigeon genome: their large population size increased both neutral genetic diversity and the efficacy of selection, but linkage between sites, particularly in genomic regions with lower recombination rates, acted to reduce genetic diversity and the efficacy of selection. This conclusion is supported by studies of other birds, which have reported a correlation between recombination rate and both diversity (*25*, *26*) and the efficacy of selection (*27*–*29*). Although, it has been argued that the correlation with the efficacy of selection could be an artefact of GC-biased gene conversion (gBGC) (*30*).

Regions of the genome with higher recombination rates are expected to accumulate GC substitutions faster as a result of gBGC. gBGC promotes the fixation of A/T to G/C mutations and the loss of G/C to A/T mutations by preferentially replacing A/T bases with G/C bases when recombination occurs at a heterozygous locus (*31*). gBGC is predicted to have a greater influence in larger populations (*32*). We observe a higher GC-content in high-recombination regions of both species’ genomes (fig. S5), which indicates a long-term influence of gBGC. We also observe a higher rate of A/T to G/C substitution and a lower rate of G/C to A/T substitution in passenger pigeons than in band-tailed pigeons, which suggests a greater influence of gBGC in passenger pigeons (Fig. 4A,B).

**Fig. 4.**
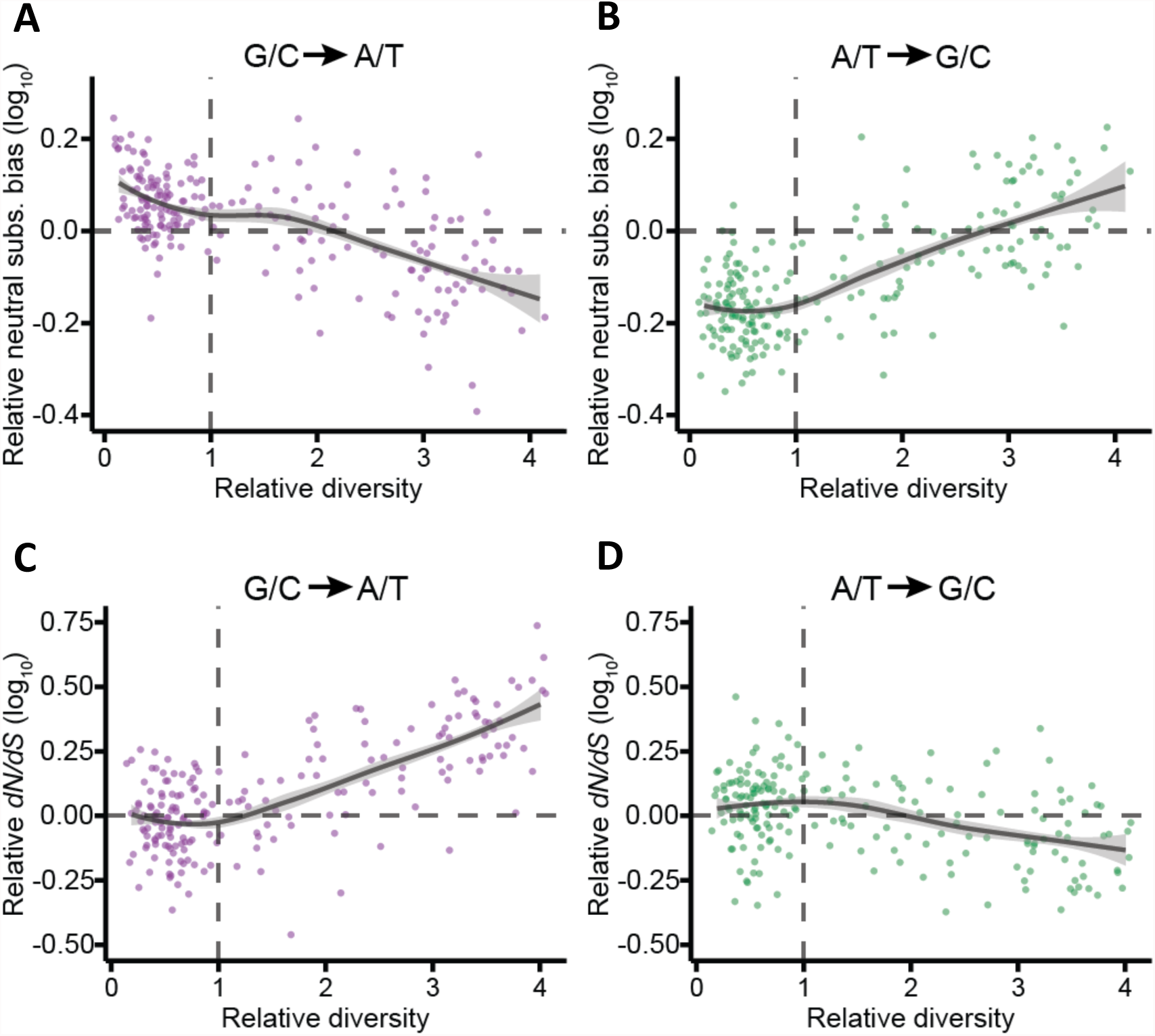
Patterns of substitution for nucleotide base changes that are opposed (A, C) and promoted (B, D) by gBGC. (**A**) The rate of G/C to A/T substitution relative to G/C to G/C substitution in passenger pigeons, divided by the same parameter in band-tailed pigeons. (**B**) The rate of A/T to G/C substitution relative to A/T to A/T substitution in passenger pigeons lineage, divided by the same parameter in band-tailed pigeons. (**C**) *dN*/*dS* for G/C to A/T mutations in passenger pigeons, divided by the same parameter in band-tailed pigeons. (**D**) *dN*/*dS* for A/T to G/C mutations in passenger pigeons, divided by the same parameter in band-tailed pigeons. All estimates are for 5 Mb windows across the genome, and are plotted on a log_10_-scale against diversity in passenger pigeons relative to band-tailed pigeons. Trend lines were estimated using the ‘stat_smooth’ function in *ggplot2* (method = ‘loess’) in *R*. Shading reflects 95% confidence limits around the trend lines.

The opposition of gBGC by selection, for example in purging deleterious G/C mutations, could therefore create the appearance of greater efficacy of selection in passenger pigeons. The impact of gBGC on signals of selection can be seen most clearly in a higher rate of nonsynonymous substitution relative to synonymous substitution (*dN*/*dS*) for substitutions opposed by gBGC and a lower *dN*/*dS* for substitutions promoted by gBGC in high-diversity regions of the passenger pigeon genome (Fig. 4C,D and fig. S6). We also find that it influences *ω_a_* and *pN*/*pS* (fig. S7 and S8). To test whether our inference of more efficient selection in passenger pigeons is an artefact of gBGC, we estimated *ω*_a_ and *pN*/*pS* separately for G/C to G/C and A/T to A/T mutations, which are unaffected by gBGC. For these mutations, we again observed higher *ω_a_* and lower *pN*/*pS* in passenger pigeons than in band-tailed pigeons (figs. S7 and S8), confirming that selection was genuinely more efficient. However, when comparing high- and low-diversity regions of the passenger pigeon genome, we only observe a difference in *pN*/*pS*. This indicates that differences in *ω_a_* across the passenger pigeon genome were driven by gBGC.

A previous study suggested that passenger pigeons’ low genetic diversity was the result of drastic population fluctuations driven by resource availability (*4*). This conclusion was based on Pairwise Sequentially Markovian Coalescent (PSMC) analyses (*33*) of the nuclear genome. In contrast, our analyses reveal both population stability preceding the species’ extinction and a surprisingly pervasive influence of natural selection. Moreover, the extent of the influence of selection across the passenger pigeon genome means that analyses that are based on genome-wide diversity, such as PSMC, are unlikely to reliably inform us of demographic history (*34*) (SM2.5). Our results therefore undermine the argument that natural demographic fluctuations contributed to the passenger pigeon’s extinction, and instead suggest that natural selection could have played a role: following the onset of the commercial harvest, low genetic diversity may have made passenger pigeons less able to respond to new selective pressures, and while the species benefitted from higher rates of adaptive evolution and efficient purifying selection, previously adaptive traits may have made it more difficult for passenger pigeons to survive in smaller numbers (*2*).

## Acknowledgments

We thank L. Shiue, S. Weber, J. Kapp, M. Stiller, T. Kuhn, S. Wagner, and R. Shaw for assistance generating data. We thank J. Novembre for advice on analysing codon usage bias. Research was supported by the Packard Foundation, the Gordon and Betty Moore Foundation, and Revive & Restore. A.E.R.S. was funded by Ciência sem Fronteiras fellowship - CAPES, Brazil. Sequencing was supported by the Dean's Office, the Vincent J. Coates Genomics Sequencing Laboratory at UC Berkeley (Berkeley sequencing supported by NIH S10 Instrumentation Grants S10RR029668 and S10RR027303), and the Danish National Sequencing Centre in Copenhagen (sequencing supported by Lundbeck Foundation grant R52-5062).

The sequence data generated in this study are archived in the relevant NCBI databases: the band-tailed pigeon assembly and RNA-seq reads used for its annotation can be found in Bioproject PRJNA308039 and reads from passenger pigeon samples in PRJNA381231 (with accessions provided in the supplementary materials).

## Author contributions

B.S. conceived and designed the study with critical input from G.G.R.M, A.E.R.S, R.E.G, and R.B.C-D.; B.S., T.L.F, and B.J.N. led sample collection; A.J.B., A.D., J.R.D., A.G.S., K.S., G.S., M.T.P.G., and M.P. provided samples; A.E.R.S., T.L.F., B.L., B.J.N, and R.R.DaF performed DNA extraction and library preparation experiments; A.E.R.S and P.D.H performed mitochondrial genome assembly and analyses; A.E.R.S, N.K.S, E.S.R, J.A.C., S.H.V., and P.D.H. performed nuclear genome assembly and analyses; G.G.R.M. designed and performed selection analyses; B.S., G.G.R.M, A.E.R.S, and R.E.G. wrote the paper; and all authors contributed to editing the manuscript.

